# Honey bee workers reared in a neonicotinoid contaminated in-hive environment

**DOI:** 10.1101/420919

**Authors:** Richard Odemer, Franziska Odemer

**Affiliations:** University of Hohenheim, Apicultural State Institute, 70593 Stuttgart, Germany

**Keywords:** neonicotinoid pesticide, clothianidin, honey bees, brood, risk assessment, field conditions, test system, terrestrial ecotoxicology

## Abstract

With the currently updated risk assessment of three neonicotinoid pesticides, the European Food Safety Authority has confirmed that different applications of these substances represent a risk to wild and managed bees and their use was therefore severely restricted. However, to close further gaps in knowledge, this experiment covers exposure of honey bee worker brood reared in a neonicotinoid contaminated in-hive environment with focus on the individual. In a worst case scenario, mini-hives were fed chronically with a sublethal concentration of clothianidin (15 µg/kg), which is highly toxic to bees already in small amounts. Freshly hatched workers from these colonies were subsequently marked and introduced into non-contaminated colonies, where their lifespan and behavior was monitored. Nineteen days after exposure, clothianidin treated bees had no reduced lifespan or showed any signs of behavioral impairment when compared to the control, demonstrating that social buffering is not a simple substitution of dead bees by rearing more brood. Our results suggest that the social environment plays a crucial role for the individual in terms of “superorganism resilience”. These findings are discussed in context with the current use of lower tier test systems in risk assessment and contrary results obtained from laboratory experiments.

**HIGHLIGHTS:**

- Sublethal clothianidin treatment did not affect lifespan nor behavior of workers.
- Effects on individual bees reared within a mini-hive are translatable to full-sized colonies.
- “Superorganism resilience” is not a simple substitution of dead bees by rearing more brood.
- Laboratory testing in the risk assessment of plant protection products bears severe weaknesses.

## 1 INTRODUCTION

Since 2013 the use of three neonicotinoid pesticides has been restricted to non-flowering crops across the EU, over concerns they can do considerable harm to managed and wild bees (EFSA 2013). The current European Commission ban expected to come into force by the end of 2018 goes much further, prohibiting their outdoor use entirely. The decision was backed up by evaluating nearly 1,600 documents, including laboratory, semi-field and field studies with honey bees, bumblebees and solitary bees published by scientists, regulatory authorities and the agrochemical industry (EFSA 2018). Demonstrating an impressive amount of research, most of these studies deal with the European honey bee (*Apis mellifera* L.) as a model species. Many laboratory tests provide evidence that even sublethal doses of neonicotinoids can affect both, behavior and physiological processes within the insect.

In particular, social behavior (Forfert & Moritz 2017; Tison et al. 2016) and the reduction of bees’ immunocompetence (Brandt et al., 2016), but also impaired navigation (Fischer et al. 2014) and compromised learning and memory functions (Tison et al. 2017) of individual bees to name but a few. Moreover, a wide range of pesticide residues from agricultural practice can be found in hive environments (Mullin et al. 2010) possibly acting as background noise.

The different exposures experienced by individual bees (nectar, pollen or other sources) causes a distribution of individual effects, ranging from mild sublethal impairment to death. More importantly, these individual effects may translate into effects on colony-level functions and should therefore be investigated with regard to such (Sponsler & Johnson 2017).

For honey bees it has become obvious that within their hive entity, a colony is able to buffer environmental impairments to a certain extent. However, the mechanisms remain not fully understood until today (Straub et al. 2015). At present, the risk assessment of plant protection products on honey bees favors a tiered approach using a screening procedure, i.e., Tier I testing, to rapidly determine which pesticides are expected to pose a minimal risk, indicating that further analysis is not necessary. This approach is exclusively based on toxicity data from studies conducted with individual bees in the laboratory (Rortais et al. 2017). Even though laboratory experiments may present clear advantages towards semi-field and field studies in terms of comparability, degree of standardization and of course cost intensity, they bear the risk of turning a blind eye to possible sublethal effects with more or less fatal consequences for the colony (Williams et al. 2013).

Common endpoints of acute toxicity are the lethal dose (LD_50_) or lethal concentration (LC_50_) that causes death (resulting from a single or limited exposure) in 50 percent of the treated individuals. The LD_50_ is generally expressed as the dose in milligrams (mg) of active ingredient per kilogram (kg) of body weight. LC_50_ is often expressed as mg of active ingredient per volume (e.g., liter (L)) of medium (i.e., air or water) the organism is exposed to. Chemicals are considered highly toxic when the LD_50_/LC_50_ is small and practically non-toxic when the value is large. However, the LD_50_/LC_50_ does not reflect any effects from long-term exposure (i.e., overwintering, behavior or reproductive success) that may occur at levels below those that cause death (Pinna 2017).

For the neonicotinoid clothianidin, Laurino et al. (2013) reported a LD_50_ of 3.54 ng/bee (24 h) and a similar result was reported from EFSA (2013) with 3.79 ng/bee (24 h). However, more recent studies showed a seven-fold lower susceptibility with 26.9 ng/bee (48 h) (Alkassab & Kirchner 2016) and 25.4 ng/bee (48 h) (Yao et al. 2018), demonstrating discrepancy and inaccuracy for this endpoint and potentially even for the test system(s).

With our approach, we wanted to (i) bridge four traditional laboratory tests frequently in use for the testing of plant protection products by Contract Research Organizations (CRO’s) issued by the OECD. At the same time this approach should (ii) represent a more realistic scenario under which honey bees are exposed to such chemicals under a reduced cost and time scope of service. Therefore, as a first step, a whole colony setup was employed by using fully intact mini-hives to evaluate lethal and sublethal effects of the highly toxic neonicotinoid clothianidin. A possible long-term impact on adult worker bees developed from larvae reared under exposure stress was evaluated and put together in context with the current procedures of laboratory testing in honey bee risk assessment.

## 2 MATERIALS & METHODS

### 2.1 Experimental setup

This experiment was performed in August and September of 2013 with the “Kieler mating nuc” system (KMN), a Styrofoam box with four top-bars, a strip of a beeswax foundation attached to it and equipped with a feeder (Fig. 1) (Odemer et al. 2018). Twelve mini-hives were established with approximately 800 bees originating from brood frames of two donor colonies, treated against Varroosis and proven to be free of *Nosema* spores (Fries et al. 2013) four weeks before the experimental start. Subsequently, unmated sister queens were introduced to the mini-hives and they were set up at a protected apiary of the institute for mating after one night in a dark and chilled room. Within a period of five weeks, all of the established hives showed several brood stages (eggs, larvae and patches of sealed brood) and freshly built wax combs as a sign of successful mating.

**Fig. 1.**
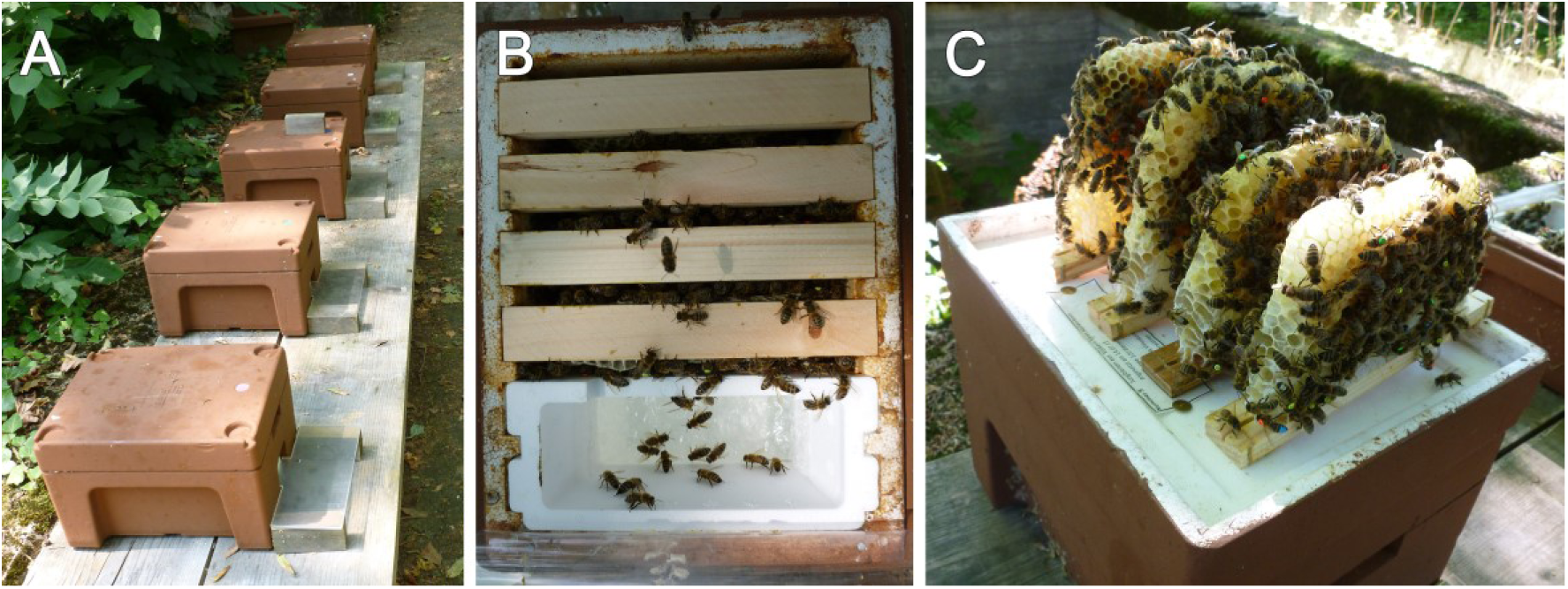
**A:** Mini-hive setup of one treatment group including five hives during the neonicotinoid exposure. **B:** top view of a mini-hive with four top-bars and a feeding device, providing space for an intact honey bee colony. **C:** for the mortality assessment, all four combs were removed and both sides of each top-bar with attached bees was photographed]

### 2.2 Experimental field site and weather conditions

The experiment was conducted at the Apicultural State Institute in Stuttgart-Hohenheim (48°42’31.8“N 9°12’38.2”E). Within the closer range of approximately 250 m, no other honey bee colonies were present. In the wider range (> 250 m), other experimental hives as well as observation hives were placed, to provide enough drones for mating. At the time present, main natural food source from local flora mainly consisted of nectar and honeydew from *Tilia spp.*.

The average temperature during the experiment was 22.5 °C with a precipitation of 101.6 L/m^2^. Overall, good weather conditions prevailed to perform foraging flights (DWD 2013).

### 2.3 Clothianidin treatment

As a nitro-substituted neonicotinoid, clothianidin is of high toxicity to honey bees (Iwasa et al. 2004). The oral LC_50_ was calculated to be 37 µg/kg or expressed as LD_50_: 3.7 ng/bee, respectively with a NOEL (No Observed Effect Level) of 20 µg/kg (Würfel 2008, Alkassab & Kirchner 2016).

For the application of clothianidin a dry compound was used (99 % purity, Dr. Ehrenstorfer GmbH), sonicated in pure water for a stock solution. The amount of stock solution was calculated for a final concentration of 15 µg/kg in sucrose feeding syrup (Apiinvert, Südzucker GmbH), which was considered to be below an acute toxic concentration (Alkassab & Kirchner 2016). The same amount of pure water without clothianidin was used for the control treatment.

### 2.4 Treatment groups

Ten of the 12 established mini-hives were split randomly into two groups of five (control and treatment). The control received feeding syrup - free of any pesticide (Tab. 1) - while the treatment group was chronically fed for 26 consecutive days with 1.68 kg feeding syrup containing a concentration of 15 µg clothianidin/kg, corresponding to a total amount of 25.2 µg clothianidin/mini-hive.

One sealed brood comb with bees about to hatch was removed from all mini-hives of both treatments and put together group wise (5 combs each) in an incubator for 24 hours. Afterwards, 100 freshly hatched bees per treatment were collected at random and individually labelled with a colored and numbered opalith plate on the thorax. In addition we marked the dorsal side of the abdomen with a hive specific color in order to identify drifting bees that enter neighboring colonies. The bees were then introduced into the two remaining mini-hives (Col1 and Col2, from the initial 12) divided in 50 treated and 50 untreated bees per hive. Subsequently, the mortality and behavioral abnormalities were recorded for a period of 19 days. See Fig. 2 for a detailed experimental setup diagram.

**Fig. 2.**
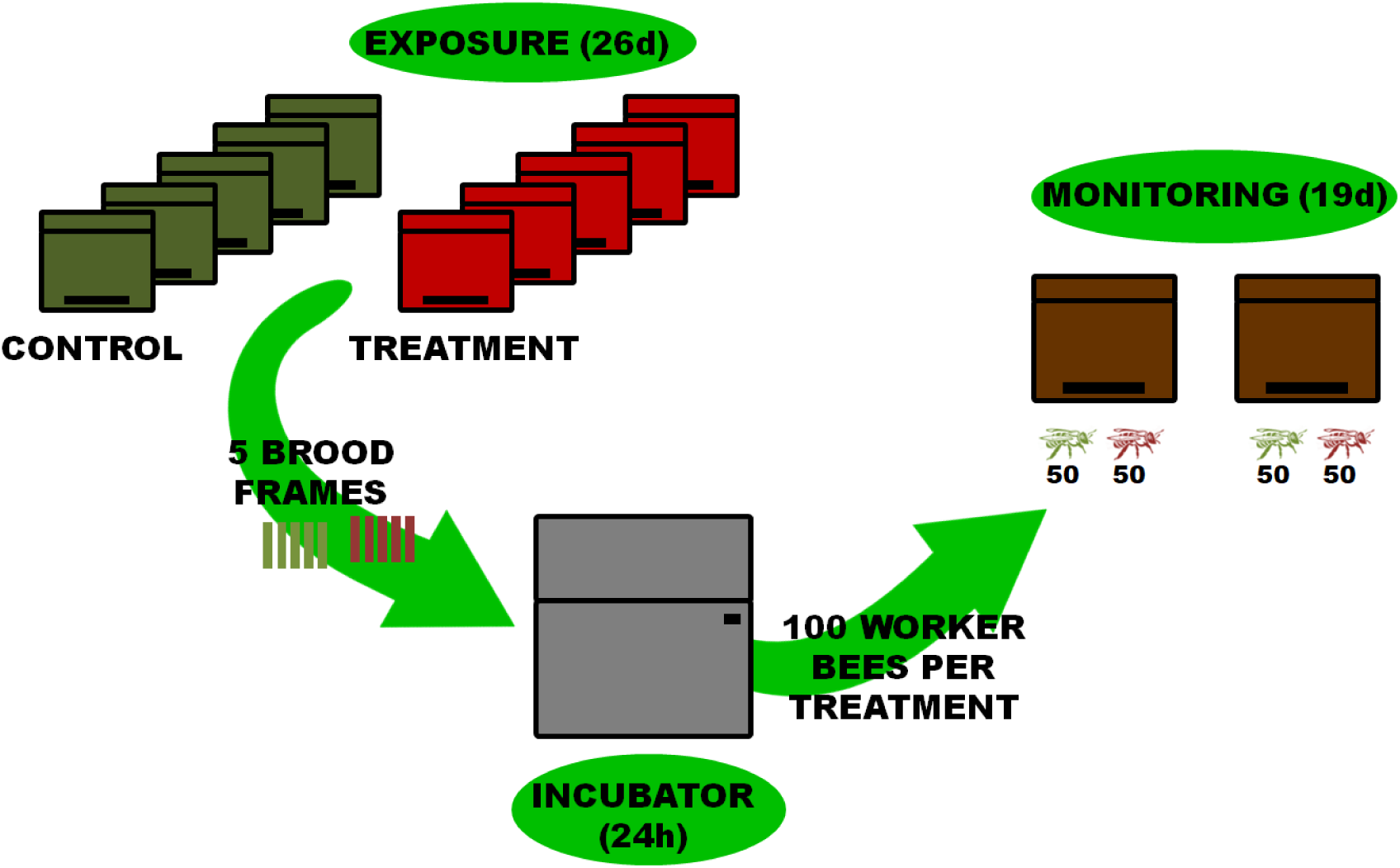
Diagram of the experimental setup illustrating the exposure and monitoring period. In the treatment, five mini-hives were exposed to clothianidin for 26 days. Subsequently, one ready-to-hatch brood frame per mini-hive was removed and placed together group wise in an incubator for 24 h. After hatching, a total of 100 worker bees per treatment were marked and introduced into two neutral mini-hives. The bees were further divided in 50 treated and 50 untreated bees per hive and monitored for 19 d.]

### 2.5 Mortality assessments

The monitoring started 24 hours after the bees’ introduction for a period of 19 days. The assessment included a daily mortality check, for which all combs including the inside of the hive were photographed for the later on counting of the marked bees on a computer screen. The pictures were taken outside the foraging activity, early in the morning. The overall recovery rate is also shown in 3.1.

### 2.6 Behavioral abnormalities

Behavioral abnormalities were recorded daily at the same time as the mortality check, quantitatively observed according to the following categories adopted from TG 245 (OECD, 2017): m = moribund (bees cannot walk and show only very feeble movements of legs and antennae, only weak response to stimulation; e.g. light or blowing; bees may recover but usually die),

a = affected (bees still upright and attempting to walk but showing signs of reduced coordination; hyperactivity; aggressiveness; increased self-cleaning behavior; rotations; shivering),

c = cramps (bees contracting abdomen or entire body),

ap = apathy (bees show only low or delayed reactions to stimulation e.g. light or puff of air; bees are sitting motionless in the unit).

v = vomiting

### 2.7 Residue analysis

Ahead of the experiment, a sample of the stock solution and the treated feeding syrup was collected. At day 18, pooled samples of pollen (bee bread) and stored food of both groups were collected from in-hive storage cells. All samples were analyzed using GC-MS and/or LC-MS/MS following acetonitrile extraction/partitioning and clean-up by dispersive Solid Phase Extraction (SPE) - QuEChERS-method; German version EN 15662:2009 in certified labs (feeding syrup and food: eurofins Dr. Specht Labs Hamburg, LOQ 3 µg/kg; pollen: LUFA Speyer, LOQ 0.3 µg/kg).

### 2.8 Statistical analysis

We evaluated the mortality data with a Kaplan-Meier-Survival analysis. Survivorship between control and treatment was compared pairwise and tested for significance with Log-Rank Tests (Cox-Mantel) followed by a Bonferroni correction. Workers which were collected at the end of the experiment were considered censored, equal to those observed but not collected on the last day of the experiment. Possible inter-colony effects were evaluated with a Cox proportional hazards model as a covariate. All tests were performed using RStudio (R Core Team, 2018) and a significance level of α=0.05.

## 3 RESULTS

### 3.1 Recovery rate of introduced bees

The recovery rate was calculated by the number of bees that could be rediscovered 24 h after the introduction of 100 particularly treated worker bees per mini-hive. The high recovery rates in all groups ranging from 99 to 100 % indicate that the prior treatment (feeding of clothianidin and hatching in the incubator) did not have an acute negative impact.

### 3.2 Residue analysis

The laboratory analysis of the stock solution and the feeding syrup verified an intended clothianidin concentration in the feeding syrup of 15 µg/kg. Additionally, we found measurable residues ranging from ∼2 to 6 µg/kg in stored food and pollen of the five clothianidin treated mini-hives. We could also confirm that the untreated controls were free of clothianidin residues (Tab. 1).

**Tab. 1.** Residue analysis of control and clothianidin treated feeding syrup prior to observation period. Pooled food and pollen from storage combs of all control and clothianidin treated mini-hives after 18 days (LC-MS/MS, LOQ: 3 µg/kg for food, 0.3 µg/kg for pollen).

|  | Control | clothianidin |
| --- | --- | --- |
| Stock solution | - | 15 mg/kg |
| Feeding syrup | 0 µg/kg | 15 µg/kg |
| Stored food | 0 µg/kg | 6 µg/kg |
| Stored pollen | < 0.3 µg/kg | 1.79 µg/kg |

### 3.3 Mortality of worker bees

The Kaplan-Meier-Survival analysis of both groups showed a significant difference indicating a lower mortality of the clothianidin treated bees when compared to the control group (p=0.043) (Fig. 3). Therefore, a Cox proportional hazards model was applied to determine the hazard ratio (HR) displayed as forest plot (Fig. 4). With a HR of 0.7 the clothianidin treated bees did not have a reduced risk of dying when compared to the control (p=0.062). In addition, the two colonies (Col1 and Col2) were compared to display possible intercolony effects. However, with a HR of 1.3 bees in Col2 did not have a higher risk of dying when compared to Col1 (p=0.158).

**Fig. 3.**
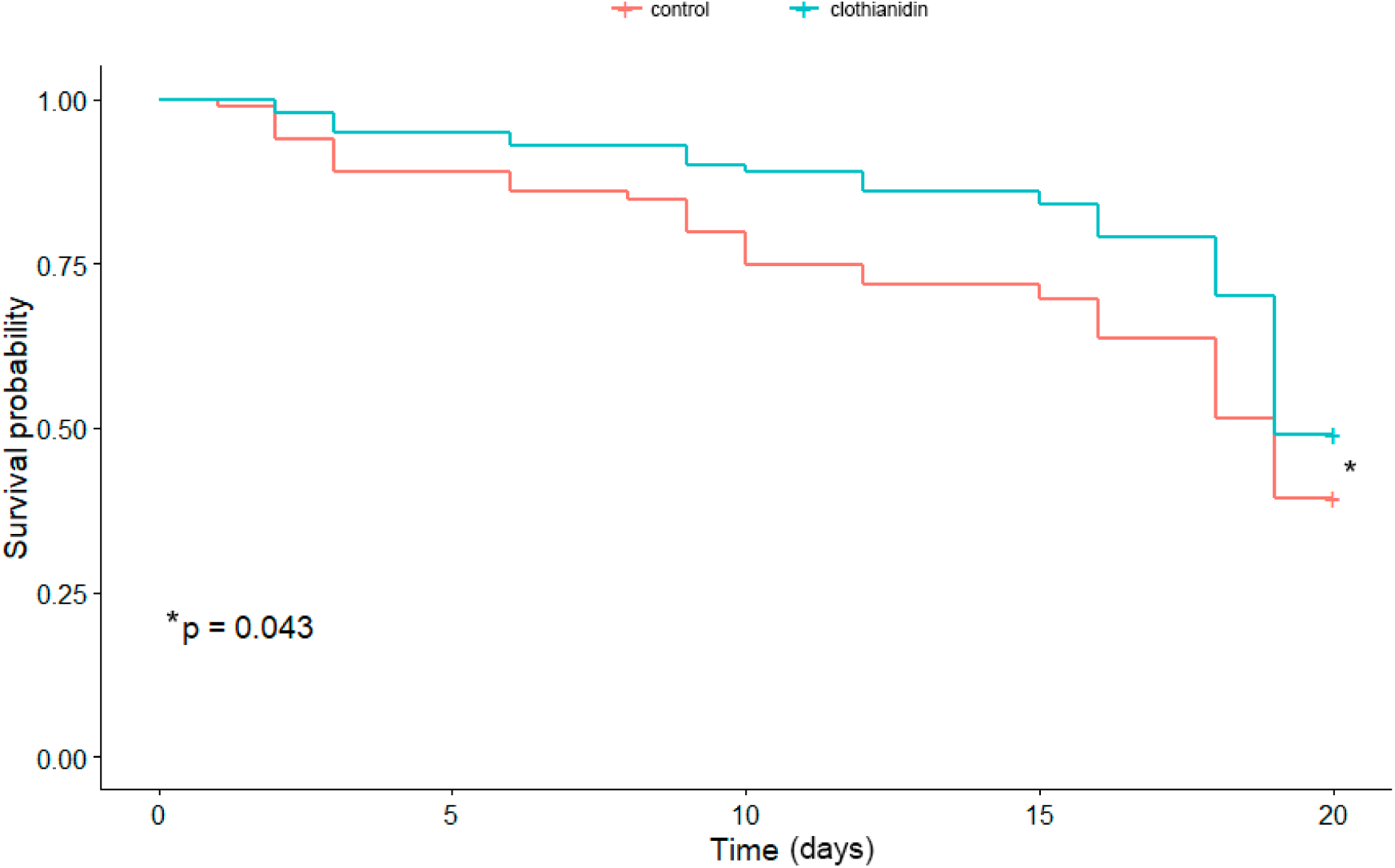
Both treatment groups were compared with a Kaplan-Meier-Survival analysis. A post-hoc Log-Rank test (Cox-Mantel) revealed significant differences between those groups (Log-Rank p=0.043). Asterisk indicates statistically significantly lower mortality when compared to the control group]

**Fig. 4.**
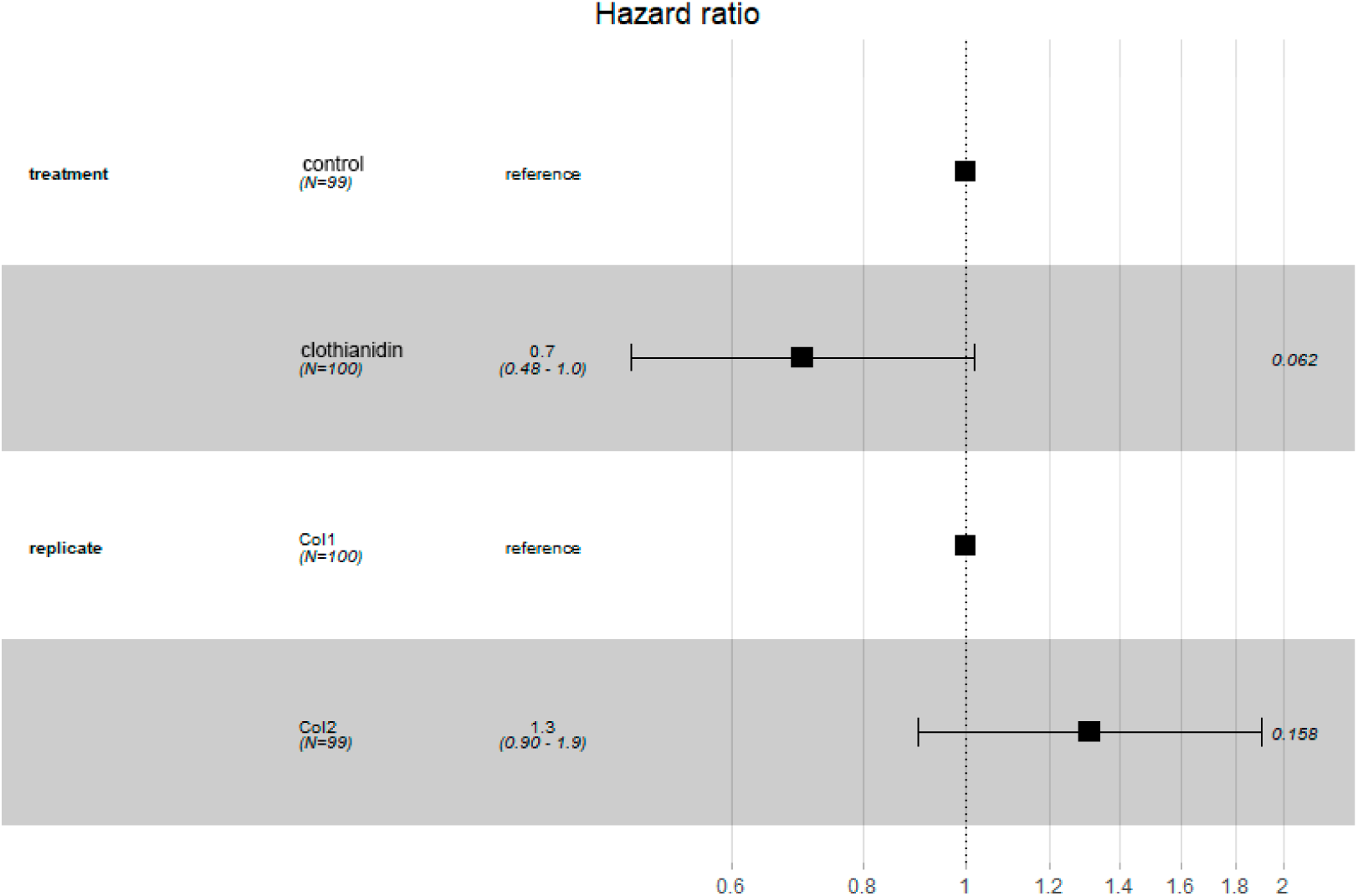
Both treatment groups and the two colonies (Col1 and Col2) were additionally compared with a Cox proportional hazards model to determine the hazard ratio (HR) displayed as forest plot. With a HR of 0.7 the clothianidin treated bees did not have a reduced risk of dying when compared to the control (p=0.062). And with a HR of 1.3 bees in Col2 did not have a higher risk of dying when compared to Col1 (p=0.158)]

### 3.1 Behavioral abnormalities

No behavioral abnormalities according to the categories in 2.6 could be noticed. Neither control nor clothianidin treated bees showed visible signs of sublethal impairment.

## 4 DISCUSSION

We found that honey bee workers previously exposed to clothianidin in their most sensitive stage of development (larvae, pupae) did not live shorter nor showed significant abnormal behavior when monitored further after reaching adulthood. This is one of the first studies that incorporated a realistic sublethal worst-case exposure scenario into laboratory testing procedures. With our results, we provide complementary evidence for “superorganism resilience” or social buffering capacity generated by intact honey bee colonies to withstand environmental stressors to several degrees (Straub et al. 2015). Even though many studies in the past decade found detrimental effects related to neonicotinoid exposure in terms of navigation (Henry et al. 2012; Fischer et al. 2014), learning ability (Decourtye et al. 2003), memory functions (Williamson & Wright 2013; Tison et al. 2017), synergistic response to pathogens (Doublet et al. 2015; Coulon et al. 2017), immunocompetence (Di Prisco et al. 2013; Brandt et al. 2016) and others, most of these findings were the result of studies focused on single bees extracted from their colony (Lundin et al. 2015). In addition, unrealistic exposure scenarios were employed, where doses or concentrations indeed were sublethal but in many cases exceeded residue levels found in practice (Blaquiere et al. 2012; Rolke et al. 2016; Mitchell et al. 2017).

Recent publications have shown honey bees that were exposed to neonicotinoid pesticides under realistic field conditions at the colony level were not adversely affected in general (Solomon & Stevenson 2017). In addition, Siede et al. (2018) found that neither colony health, bee mortality, overwintering success nor hive weights were impaired by a sublethal clothianidin treatment. However, *Varroa destructor* had a significant negative impact on colony strength but no correlations could be show for a synergism between the parasite and the pesticide treatment. This is in line with two previous studies that we have conducted under comparable conditions. In Retschnig et al. (2015) we were able to show that environmental factors such as the study location may play a crucial role in pesticide susceptibility, corroborated by Woodcock et al. (2017) with a similar conclusion. And in Odemer et al. (2018) treatment effects from clothianidin as well as a potential synergism with the intracellular gut parasites *Nosema spp.* failed to appear. Also for the neonicotinoid thiacloprid frequently used in flowering crops attractive to pollinators, diverse impairment as a result from laboratory or semi-field studies could not be observed (Tison et al. 2016, 2017; Forfert & Moritz 2017). Under field conditions however, monitored full sized colonies did not show negative effects on overwintering success nor on colony development (Odemer & Rosenkranz 2018), closely in line with the results from Siede et al. (2017) and large scale monitoring projects such as the German bee monitoring DEBIMO (Genersch et al. 2010) or the pan-European EPILOBEE (Jacques et al. 2017). As a social entity, fundamental performance parameters of a honey bee colony are obviously isolated from the fate of individuals suffering death or other negative impact by a highly complex interaction of compensatory mechanisms (Berenbaum & Johnson 2015, Sponsler & Johnson 2017, Manjon et al. 2018). Among others, performance parameters such as overwintering success, population dynamics and honey yield are endpoints of higher tier studies in the current honey bee risk assessment (Rortais et al. 2017). One aim of our approach was to observe effects of toxic exposure after all the mechanisms of “social buffering” have played their role (Sponsler & Johnson 2017). We could demonstrate that even in a smaller scale, it was possible to create a realistic exposure scenario with fully intact honey bee colonies as a way of supplementing the existing laboratory test systems.

The current OECD test guidelines and guidance documents including the acute and chronic oral toxicity test on adult workers (OECD 1998, 2017) and the larval toxicity test acute and repeated exposure (OECD 2013, 2016) represent full-featured laboratory test systems for risk assessment purposes. Honey bee workers and larvae are reared under artificial conditions fed with a synthetic diet in an incubator removed from their natural hive environment. Although intended for screening purposes only, these test systems represent the first step of an important decision process if further testing is necessary or not. The level of standardization is comparatively high, environmental factors are entirely controllable for which an increased comparability is given. Feeding and frequent mortality assessments are time and care consuming, as negligence immediately results in dead test subjects. In addition, none of these tests are using a diet appropriate to the test subject, e.g. adult honey bees are reared without a protein source and larvae are fed an artificial diet containing commercially available royal jelly (RJ).

Thereby, pollen is known to be potentially involved in bee health and defense response (Di Pasquale et al. 2013) and a deficit can thus be responsible for higher mortality rates in cage experiments (Frias et al. 2015). Rittschof et al. (2015) found that the pre-adult environment of workers had a significant influence on the resilience to immune challenges in the form of pesticide exposure. Even though the molecular mechanisms remain yet unknown, a possible candidate system in pesticide response is assumed. Cytochrome P450 enzymes are adjusted to nutritional ingredients in honey bee adults, and potentially in larvae (Cameron et al. 2013; Mao et al. 2016). Highly relevant for detoxification (Ferguson & Tyndale 2011), CYP9Q3 was recently identified to be substantially involved in the process (Manjon et al. 2018). It is suggested that the larval diet may have a long-term impact on P450 levels in adult bees resulting in defense against pesticide toxication (Rittschof et al. 2015). This developmental step is excluded in both OECD larval guidelines with unknown consequences for the test results. Moreover, the study design intends to feed larvae with commercially available RJ, which is most commonly imported from China (Cao et al. 2016). As for retailers it is not required to provide a certificate of analysis with their product, while residue studies have revealed that commercial RJ can be contaminated with antibiotics such as chloramphenicol (Calvarese at al. 2006, Veterinary Residues Committee 2007), tetracyclines and sulfonamides (Gunes et al. 2009) to treat against bacterial brood diseases, for instance American and European foulbrood (Rama et al. 2015). Prophylactic residue analyses are time consuming and expensive, hence most CRO’s do their testing without a safety feature of this sort, at the risk of contamination. In addition, recent studies have shown that there are substantial disparities in the nutritional composition of RJ and worker jelly (WJ). Wang et al. (2018) found a different proportion of sugars, protein and trace elements varying also on a daily basis. Moreover, Buttstedt et al. (2018) discovered the composition of protein controlling viscosity in RJ ensures that the queen larva vertically attached to its cell doesn’t fall out. This suggests that such differences have a potential impact on the development and physiology of artificially reared larvae according to OECD TG 237 and GD 239 (OECD 2013, 2016). Therefore the outcome of these studies and what they represent may be questioned.

Our approach, in contrast, uses the colony to provide environmental conditions and feeding, appropriate for either larvae or adult workers on a relatively low cost level. It is neither mandatory to mix artificial diet nor feeding solutions to meet the minimal requirements for the subjects to survive nor is there a risk of contamination from external sources. The colony makes the optimal choice for each individual and each developmental stage, representing the most realistic condition for these tasks (Seeley, 2009). Even if for now, only mortality was in focus of our test system, we could show that handling and practicability of the test subjects in terms of application, maintenance, photographic assessments and evaluation of results was well suited for an initial trial. Once enhanced with the right technique, i.e. photographic assessment software (Höferlin et al. 2013), RFID entrance readers for incoming and outgoing bees (de Souza et al. 2018), radio controlled scales (Meikle et al. 2018), our test system is capable of providing precise results for endpoints such as daily food uptake, brood termination rate, flight activity and others suitable for augmenting good laboratory practice (GLP) testing.

## Conclusion

Our test system was able to generate an outcome comparable to larger scale field studies, demonstrating for the first time that effects on individual bees reared within a mini-hive are translatable to full-sized colonies. We provide evidence that the “superorganism resilience” is not a simple substitution of dead bees by rearing more brood, but rather a lower vulnerability of the individual within its social environment.

Furthermore, we revealed that laboratory testing in the risk assessment of plant protection products bears severe weaknesses which may result in misjudgments and ultimately in the admission of harmful products. Hence, not only pollinators but whole ecosystems are put at risk. We therefore suggest that such test systems need revision having regard to the biological relevance of the whole colony in addition to the individual. With further enhancement, the mini-hive would make an appropriate test system for initial screening studies of plant protection products supplementing or replacing cage experiments in the laboratory.

## ACKNOWLEDGEMENTS

We appreciate the support of Peter Rosenkranz by providing the possibility to conduct the experiment within the facilities of the Apicultural State Institute in Hohenheim.

This research did not receive any specific grant from funding agencies in the public, commercial, or not-for-profit sectors.

## Ethical approval

This article does not contain any studies with human participants or animals performed by any of the authors.

## REFERENCES

Alkassab AT, Kirchner WH (2016) Impacts of chronic sublethal exposure to clothianidin on winter honeybees. Ecotoxicology 25:1000–1010. doi:10.1007/s10646-016-1657-3

Berenbaum MR, Johnson RM (2015) Xenobiotic detoxification pathways in honey bees. Curr Opin Insect Sci 10:51–58. doi: 10.1016/j.cois.2015.03.005

Blacquière T, Smagghe G, Van Gestel CAM, Mommaerts V (2012) Neonicotinoids in bees: A review on concentrations, side-effects and risk assessment. Ecotoxicology 21:973–992. doi: 10.1007/s10646-012-0863-x

Brandt A, Gorenflo A, Siede R, Meixner M, Büchler R, B??chler R (2016) The neonicotinoids thiacloprid, imidacloprid, and clothianidin affect the immunocompetence of honey bees (Apis mellifera L.). J Insect Physiol 86:40–47. doi: 10.1016/j.jinsphys.2016.01.001

Buttstedt A, Muresan CI, Lilie H, Hause G, Ihling CH, Schulze S-H, Pietzsch M, Moritz RFA (2018) How Honeybees Defy Gravity with Royal Jelly to Raise Queens. Curr Biol 28:1095–1100.e3. doi: 10.1016/j.cub.2018.02.022

Calvarese S, Forti AF, Scortichini G, Diletti G (2006) Chloramphenicol in royal jelly: analytical aspects and occurrence in Italian imports. Apidologie 37:673–678. doi: 10.1051/apido:2006042

Cao L-F, Zheng H-Q, Pirk CWW, Hu F-L, Xu Z-W (2016) High Royal Jelly-Producing Honeybees (Apis mellifera ligustica) (Hymenoptera: Apidae) in China. J Econ Entomol 109:510–514. doi: 10.1093/jee/tow013

Cameron RC, Duncan EJ, Dearden PK (2013) Biased gene expression in early honeybee larval development. BMC Genomics 14:903. doi: 10.1186/1471-2164-14-903

Coulon M, Schurr F, Martel A-C, Cougoule N, Bégaud A, Mangoni P, Dalmon A, Alaux C, Le Conte Y, Thiéry R, Ribière-Chabert M, Dubois E (2017) Metabolisation of thiamethoxam (a neonicotinoid pesticide) and interaction with the Chronic bee paralysis virus in honeybees. Pestic Biochem Physiol 0–1. doi: 10.1016/j.pestbp.2017.10.009

Dai P, Jack CJ, Mortensen AN, Bustamante TA, Bloomquist JR, Ellis JD (2018) Chronic toxicity of clothianidin, imidacloprid, chlorpyrifos, and dimethoate to Apis mellifera L. larvae reared in vitro. Pest Manag Sci. doi: 10.1002/ps.5124

Decourtye A, Lacassie E, Pham-Delégue MH (2003) Learning performances of honeybees (Apis mellifera L) are differentially affected by imidacloprid according to the season. Pest Manag Sci 59:269–278. doi: 10.1002/ps.631

de Souza P, Marendy P, Barbosa K, Budi S, Hirsch P, Nikolic N, Gunthorpe T, Pessin G, Davie A (2018) Low-Cost Electronic Tagging System for Bee Monitoring. Sensors 18:2124. doi: 10.3390/s18072124

Deutscher Wetterdienst DWD (2013) Deutschlandwetter im Juli 2013 Sonnig, warm und trocken - ein Sommermonat wie aus dem Bilderbuch. Pressemitteilung. www.dwd.de, Accessed 26 January 2014

Di Pasquale G, Salignon M, Le Conte Y, Belzunces LP, Decourtye A, Kretzschmar A, Suchail S, Brunet J-L, Alaux C (2013) Influence of Pollen Nutrition on Honey Bee Health: Do Pollen Quality and Diversity Matter? PLoS One 8:e72016. doi: 10.1371/journal.pone.0072016

Di Prisco G, Cavaliere V, Annoscia D, Varricchio P, Caprio E, Nazzi F, Gargiulo G, Pennacchio F (2013) Neonicotinoid clothianidin adversely affects insect immunity and promotes replication of a viral pathogen in honey bees. Proc Natl Acad Sci 110:18466–18471. doi: 10.1073/pnas.1314923110

Doublet V, Labarussias M, de Miranda JR, Moritz RFA, Paxton RJ (2015) Bees under stress: Sublethal doses of a neonicotinoid pesticide and pathogens interact to elevate honey bee mortality across the life cycle. Environ Microbiol 17:969–983. doi: 10.1111/1462-2920.12426

EFSA (2013) Conclusion on the peer review of the pesticide risk assessment for bees for the active substance clothianidin. EFSA J 11:3066. doi: 10.2903/j.efsa.2013.3066

EFSA (2018) Peer review of the pesticide risk assessment for bees for the active substance clothianidin considering the uses as seed treatments and granules. EFSA J 16:4210. doi: 10.2903/j.efsa.2018.5177

EFSA (2013) COMMISSION IMPLEMENTING REGULATION (EU) No 485/2013 of 24 May 2013 amending Implementing Regulation (EU) No 540/2011, as regards the conditions of approval of the active substances clothianidin, thiamethoxam and imidacloprid, and prohibiting the use and s. Off J Eur Union L 139:12–26. doi: 10.2903/j.efsa.2013.3067.

Fischer J, Müller T, Spatz A-K, Greggers U, Grünewald B, Menzel R (2014) Neonicotinoids interfere with specific components of navigation in honeybees. PLoS One 9:e91364. doi: 10.1371/journal.pone.0091364

Forfert N, Moritz RFA (2017) Thiacloprid alters social interactions among honey bee workers (Apis mellifera). J Apic Res 56:467–474. doi: 10.1080/00218839.2017.1332542

Frias BED, Barbosa CD, Lourenço AP (2016) Pollen nutrition in honey bees (Apis mellifera): impact on adult health. Apidologie 47:15–25. doi: 10.1007/s13592-015-0373-y

Fries I, Chauzat M-P, Chen Y-P, Doublet V, Genersch E, Gisder S, Higes M, McMahon DP, Martín-Hernández R, Natsopoulou M, Paxton RJ, Tanner G, Webster TC, Williams GR (2013) Standard methods for Nosema research. J Apic Res 52:1–28. doi: 10.3896/IBRA.1.52.1.14

Genersch E, Von Der Ohe W, Kaatz H, Schroeder a, Otten C, BÃ¼chler R, Berg S, Ritter W, MÃ¼hlen W, Gisder S, Meixner M, Liebig G, Rosenkranz P (2010) The German bee monitoring project: A long term study to understand periodically high winter losses of honey bee colonies. Apidologie 41:332–352. doi: 10.1051/apido/2010014

Gunes ME, Gunes N, Cibik R (2009) Oxytetracycline and sulphonamide residues analysis of honey samples from southern Marmara region in Turkey. Bulg J Agric Sci 15:163–167.

Henry M, Beguin M, Requier F, Rollin O, Odoux J-F, Aupinel P, Aptel J, Tchamitchian S, Decourtye A (2012) A Common Pesticide Decreases Foraging Success and Survival in Honey Bees. Science (80-) 336:348–350. doi: 10.1126/science.1215039

Höferlin B, Höferlin M, Kleinhenz M, Bargen H (2013) Automatische Auswertung von Apis mellifera Wabenfotos und Brutentwicklung.

Iwasa T, Motoyama N, Ambrose JT, Roe RM (2004) Mechanism for the differential toxicity of neonicotinoid insecticides in the honey bee, Apis mellifera. Crop Prot 23:371–378. doi: 10.1016/j.cropro.2003.08.018

Jacques A, Laurent M, Ribière-Chabert M, Saussac M, Bougeard S, Budge GE, Hendrikx P, Chauzat M-P (2017) A pan-European epidemiological study reveals honey bee colony survival depends on beekeeper education and disease control. PLoS One 12:e0172591. doi: 10.1371/journal.pone.0172591

Laurino D, Manino A, Patetta A, Porporato M (2013) Toxicity of neonicotinoid insecticides on different honey bee genotypes. Bull Insectology 66:119–126.

Lundin O, Rundlöf M, Smith HG, Fries I, Bommarco R (2015) Neonicotinoid Insecticides and Their Impacts on Bees: A Systematic Review of Research Approaches and Identification of Knowledge Gaps. PLoS One 10:e0136928. doi: 10.1371/journal.pone.0136928

Manjon C, Troczka BJ, Zaworra M, Beadle K, Randall E, Hertlein G, Singh KS, Zimmer CT, Homem RA, Lueke B, Reid R, Kor L, Kohler M, Benting J, Williamson MS, Davies TGE, Field LM, Bass C, Nauen R (2018) Unravelling the Molecular Determinants of Bee Sensitivity to Neonicotinoid Insecticides. Curr Biol 1–7. doi: 10.1016/j.cub.2018.02.045

Mao W, Schuler MA, Berenbaum MR (2016) Honey constituents up-regulate detoxification and immunity genes in the western honey bee Apis mellifera. Proc Natl Acad Sci 113:E488–E488. doi: 10.1073/pnas.1525259113

Meikle WG, Holst N, Colin T, Weiss M, Carroll MJ, McFrederick QS, Barron AB (2018) Using within-day hive weight changes to measure environmental effects on honey bee colonies. PLoS One 13:e0197589. doi: 10.1371/journal.pone.0197589

Mitchell EAD, Mulhauser B, Mulot M, Mutabazi A, Glauser G, Aebi A (2017) A worldwide survey of neonicotinoids in honey. Science (80-) 358:109–111. doi: 10.1126/science.aan3684

Mullin CA, Frazier M, Frazier JL, Ashcraft S, Simonds R, Vanengelsdorp D, Pettis JS (2010) High levels of miticides and agrochemicals in North American apiaries: implications for honey bee health. PLoS One 5:e9754. doi: 10.1371/journal.pone.0009754

Odemer R, Nilles L, Linder N, Rosenkranz P (2018) Sublethal effects of clothianidin and Nosema spp. on the longevity and foraging activity of free flying honey bees. Ecotoxicology. doi: 10.1007/s10646-018-1925-5

Odemer R, Rosenkranz P (2018) Chronic exposure to a neonicotinoid pesticide and a synthetic pyrethroid in full-sized honey bee colonies. bioRxiv 293167. doi: 10.1101/293167

OECD (1998) Honeybees, Acute Oral Toxicity Test TG 213.

OECD OECD (2013) Honey bee (Apis mellifera) larval toxicity test, single exposure TG 237. 23:1–22. doi: 10.1787/9789264203785-en

OECD (2017) HONEY BEE (APIS MELLIFERA L.), CHRONIC ORAL TOXICITY TEST (10-DAY FEEDING) TG 245. 16.

OECD (2016) Honey Bee Larval Toxicity Test following Repeated Exposure GD 239. Development 33:1–16. doi: ENV/JM/MONO(2007)10

Pinna D (2017) Coping with biological growth on stone heritage objects: methods, products, applications, and perspectives, CRC Press, Canada

R Core Team (2018) R: A language and environment for statistical computing. R Foundation for Statistical Computing, Vienna, Austria.

Rama C, Rao M, Cyril L, Kumar A, Sekharan CB (2015) Quantitative Analysis of Oxytetracycline Residues in Honey by High Performance Liquid Chromatography. Int Res J Biol Sci Int Res J Biol Sci 4:2278–3202.

Retschnig G, Williams GR, Odemer R, Boltin J, Di Poto C, Mehmann MM, Retschnig P, Winiger P, Rosenkranz P, Neumann P (2015) Effects, but no interactions, of ubiquitous pesticide and parasite stressors on honey bee (Apis mellifera) lifespan and behaviour in a colony environment. Environ Microbiol 17:4322–4331. doi: 10.1111/1462-2920.12825

Rittschof CC, Coombs CB, Frazier M, Grozinger CM, Robinson GE (2015) Early-life experience affects honey bee aggression and resilience to immune challenge. Sci Rep 5:15572. doi: 10.1038/srep15572

Rolke D, Persigehl M, Peters B, Sterk G, Blenau W (2016) Large-scale monitoring of effects of clothianidin-dressed oilseed rape seeds on pollinating insects in northern Germany: residues of clothianidin in pollen, nectar and honey. Ecotoxicology 25:1691–1701. doi: 10.1007/s10646-016-1723-x

Rortais A, Arnold G, Dorne J-L, More SJ, Sperandio G, Streissl F, Szentes C, Verdonck F (2017) Risk assessment of pesticides and other stressors in bees: Principles, data gaps and perspectives from the European Food Safety Authority. Sci Total Environ 587–588:524–537. doi: 10.1016/j.scitotenv.2016.09.127

Seeley TD, (2009) The Wisdom of the Hive: the social physiology of honey bee colonies, Harvard University Press, USA

Siede R, Faust L, Meixner MD, Maus C, Grünewald B, Büchler R (2017) Performance of honey bee colonies under a long-lasting dietary exposure to sublethal concentrations of the neonicotinoid insecticide thiacloprid. Pest Manag Sci 73:1334–1344. doi: 10.1002/ps.4547

Siede R, Meixner MD, Almanza MT, Schöning R, Maus C, Büchler R (2018) A long-term field study on the effects of dietary exposure of clothianidin to varroosis-weakened honey bee colonies. Ecotoxicology. doi: 10.1007/s10646-018-1937-1

Sponsler DB, Johnson RM (2017) Mechanistic modeling of pesticide exposure: The missing keystone of honey bee toxicology. Environ Toxicol Chem 36:871–881. doi: 10.1002/etc.3661

Stephenson GL, Solomon KR (2017) Quantitative weight of evidence assessment of higher tier studies on the toxicity and risks of neonicotinoids in honeybees. 3. Clothianidin. J Toxicol Environ Heal - Part B Crit Rev 20:365–382. doi: 10.1080/10937404.2017.1388568

Straub L, Williams GR, Pettis J, Fries I, Neumann P (2015) Superorganism resilience: eusociality and susceptibility of ecosystem service providing insects to stressors. Curr Opin Insect Sci 12:109–112. doi: 10.1016/j.cois.2015.10.010

The Veterinary Residues Committee (2007) Annual Report on Surveillance for Veterinary Residues in Food in the UK 2007.

Tison L, Hahn M-L, Holtz S, Rößner A, Greggers U, Bischoff G, Menzel R (2016) Honey Bees’ Behavior Is Impaired by Chronic Exposure to the Neonicotinoid Thiacloprid in the Field. Environ Sci Technol 50:7218–7227. doi: 10.1021/acs.est.6b02658

Tison L, Holtz S, Adeoye A, Kalkan Ö, Irmisch NS, Lehmann N, Menzel R (2017) Effects of sublethal doses of thiacloprid and its formulation Calypso (r) on the learning and memory performance of honey bees. J Exp Biol 220:3695–3705. doi: 10.1242/jeb.154518

Wang Y, Ma L, Zhang W, Cui X, Wang H, Xu B (2016) Comparison of the nutrient composition of royal jelly and worker jelly of honey bees (Apis mellifera). Apidologie 47:48–56. doi: 10.1007/s13592-015-0374-x

Williams GR, Alaux C, Costa C, Csáki T, Doublet V, Eisenhardt D, Fries I, Kuhn R, McMahon DP, Medrzycki P, Murray TE, Natsopoulou ME, Neumann P, Oliver R, Paxton RJ, Pernal SF, Shutler D, Tanner G, van der Steen Jjm, Brodschneider R (2013) Standard methods for maintaining adult Apis mellifera in cages under in vitro laboratory conditions. J Apic Res 52:1–36. doi: 10.3896/IBRA.1.52.1.04

Williamson SM, Wright GA (2013) Exposure to multiple cholinergic pesticides impairs olfactory learning and memory in honeybees. J Exp Biol 216:1799–1807. doi: 10.1242/jeb.083931

Woodcock BA, Bullock JM, Shore RF, Heard MS, Pereira MG, Redhead J, Ridding L, Dean H, Sleep D, Henrys P, Peyton J, Hulmes S, Hulmes L, Sárospataki M, Saure C, Edwards M, Genersch E, Knäbe S, Pywell RF (2017) Country-specific effects of neonicotinoid pesticides on honey bees and wild bees. Science (80-) 356:1393–1395. doi: 10.1126/science.aaa1190

Würfel T (2008) Abschlussbericht Beizung und Bienenschäden.

Yao J, Zhu YC, Adamczyk J (2018) Responses of Honey Bees to Lethal and Sublethal Doses of Formulated Clothianidin Alone and Mixtures. J Econ Entomol 111:1517–1525. doi: 10.1093/jee/toy140

